# The DNA sensors AIM2 and IFI16 are NET-binding SLE autoantigens

**DOI:** 10.1101/2021.08.19.456941

**Authors:** Brendan Antiochos, Paride Fenaroli, Avi Rosenberg, Alan N. Baer, Jungsan Sohn, Jessica Li, Michelle Petri, Daniel W. Goldman, Christopher Mecoli, Livia Casciola-Rosen, Antony Rosen

**Affiliations:** Johns Hopkins University School of Medicine, Division of Rheumatology; Nephrology Unit, Parma University Hospital, Department of Medicine and Surgery, Parma, Italy; Johns Hopkins University School of Medicine, Division of Pathology; Johns Hopkins University School of Medicine, Department of Biophysics and Biophysical Chemistry

**Keywords:** Systemic Lupus Erythematosus, Neutrophil Extracellular Traps, Autoantibodies, Autoimmunity

## Abstract

Nucleic acid binding proteins are frequently targeted as autoantigens in systemic lupus erythematosus (SLE) and other interferon (IFN)-linked rheumatic diseases. The AIM-like receptors (ALRs) are IFN-inducible innate sensors that form supramolecular assemblies along double-stranded DNA of various origins. Here, we identify the ALR Absent in melanoma 2 (AIM2) as a novel autoantigen in SLE, with similar properties to the established ALR autoantigen interferon-inducible protein 16 (IFI16). Our SLE cohort revealed a frequent co-occurrence of anti-AIM2, anti-IFI16 and anti-DNA antibodies, and higher clinical measures of disease activity in patients positive for antibodies against these ALRs. We examined neutrophil extracellular traps (NETs) as DNA scaffolds on which these antigens might interact in a pro-immune context, finding that both ALRs bind NETs in vitro and in SLE renal tissues. We demonstrate that ALR binding causes NETs to resist degradation by DNase I, suggesting a mechanism whereby extracellular ALR-NET interactions may promote sustained IFN signaling. Our work suggests that extracellular ALRs bind NETs, leading to DNase resistant nucleoprotein fibers that are targeted as autoantigens in SLE.

## Introduction

Systemic lupus erythematosus (SLE) is a rheumatic disease characterized by upregulated interferon (IFN) expression and autoantibody production (1). Autoantibodies inform the identification of specific disease phenotypes and also provide insight into the mechanisms operative in rheumatic diseases (2). Many SLE autoantigens are nucleic acid binding proteins, and nucleic acid containing immune complexes are implicated in aspects of pathogenesis (3).

The AIM2-like receptors (ALRs) are a group of IFN-induced innate sensors of double-stranded (ds) DNA. AIM2 and IFI16 are the most studied members of the ALR family, which also includes IFIX and MNDA. The ALRs bind to dsDNA in a sequence-independent manner via electrostatic interactions with the dsDNA backbone, and form an oligomerized filament along areas of accessible dsDNA of any origin (4, 5). These innate sensors equip the cell with a means of identifying harmful stimuli, including viral genomes, mislocalized mitochondrial DNA, and chromosomal DNA from tumor cells. Once activated, the ALRs activate downstream innate immune signaling by type I IFN and inflammasome (IL-1/IL-18) pathways (6, 7).

Anti-IFI16 antibodies occur in both SLE and Sjogren’s Syndrome (SS), but we have previously reported that the targeted epitopes differ in these diseases (8, 9). IFI16 oligomers appear to be recognized by SS sera, suggesting that dsDNA binding may enhance its antigenicity. While AIM2 assembles similar filamentous structures on dsDNA, its status as an autoantigen has not been reported. Here, we identify AIM2 as an autoantigen in SLE (targeted in 31.3% of patients), with antibodies against AIM2, IFI16 and dsDNA being highly associated with one another. To understand why anti-ALR and anti-dsDNA antibodies might be closely co-targeted in SLE, we considered the possibility that ALRs bind to neutrophil extracellular traps (NETs) in the extracellular space. NETs are microbicidal structures consisting of protein-laden chromatin fibers generated by neutrophils in response to various stimuli (10). The NET dsDNA scaffold is a structure on which a variety of molecules interact (11), representing a platform for antigenic materials (including SLE autoantigens) to be presented to the adaptive immune system (12). We find that both AIM2 and IFI16 bind NETs in vitro and in tissues, with their binding yielding polymeric structures that confer resistance to DNase I. Together, our findings demonstrate that AIM2 and IFI16 are NET-bound autoantigens in SLE.

## Methods

### Patients

Plasma from 131 SLE patients (defined by the SLICC criteria(13)) in the Hopkins Lupus Cohort was studied for autoantibodies. Sera from 49 healthy controls was analyzed to establish a threshold for assay positivity. 133 primary Sjögren’s Syndrome (SS) patients (defined by ACR/EULAR criteria(14)) were included as disease controls. All patients and healthy controls gave informed consent for blood used in research and all work involving human subjects was approved by the Johns Hopkins Institutional Review Board. Paraffin sections from SLE renal biopsies were obtained for immunostaining and are detailed in Supplemental Table 3.

### AIM2 autoantibody assay

Full length AIM2 cDNA was subcloned into the pET28 vector (Novagen) and used to generate ^35^S-methionine labelled AIM2 protein by *in vitro* transcription and translation (IVTT) (Promega). Immunoprecipitations (IP) were performed using IVTT product diluted in Lysis Buffer (20 mM Tris pH 7.4, 150 mM NaCl, 1mM EDTA pH 7.4, 1% NP40) and 1 microliter of serum (90 minutes, 4°C). 20 microliters of Protein G Dynabeads (Thermo Fisher) were then added to each IP, and incubated for 60 minutes. Beads were magnetically isolated, washed, and boiled in gel application buffer. IP products were electrophoresed on SDS-polyacrylamide gels and visualized by fluorography. Films were scanned and AIM2 bands quantified using Quantity One software (Bio Rad). IP products were normalized to the same positive reference serum included on each gel. The cutoff for antibody positivity was set at 2 standard deviations above the mean control serum value. IFI16 antibodies were assayed by ELISA as described (15).

### NET assays

Neutrophils were isolated from healthy control PBMCs using Ficoll-Paque density gradient followed by RBC lysis using ACK buffer (Quality Biological). NET formation was induced using PMA at 100 nM for 3 hours. For immunofluorescence studies, neutrophils were plated on glass coverslips for 15 minutes prior to PMA treatment. For quantitative DNAse protection assays, NETs were induced with PMA in 96 well plates, incubated with or without purified ALRs, then treated with DNAse I at room temperature (RT) prior to incubation with 5 µM Sytox Green (Thermo Fisher) and quantification via fluorimetry using a Perkin Elmer plate reader. Experiments were performed twice.

### Immunofluorescence

Neutrophil samples were stained with anti-MPO-FITC antibody and mounted in DAPI-containing ProLong Gold Antifade Mountant (Thermo Fisher Scientific). AIM2 and IFI16 proteins were expressed, purified and fluorescently labeled as previously described (4, 5). SLE renal biopsies were stained as previously described (8) using anti-MPO rabbit polyclonal (ThermoFisher), anti-MPO mouse monoclonal (ThermoFisher), anti-IFI16 mouse monoclonal (Sigma), anti-AIM2 rabbit polyclonal (Sigma) and Hoechst 33342 (ThermoFisher). Confocal imaging was performed with a Zeiss AxioObserver with 780-Quasar confocal module.

### Statistics

Features of patients with and without AIM2 antibodies were compared using Fisher’s exact test for categorical variables and the Mann-Whitney test for continuous variables. Multivariable logistic regression was utilized to determine associations between variables. P values less than 0.05 were considered statistically significant.

## Results

### AIM2 autoantibodies are present in SLE, and frequently co-occur with anti-IFI16 and anti-dsDNA antibodies

To determine whether AIM2 was a target of the humoral immune response in SLE, we developed an IP assay to screen for anti-AIM2 antibodies. 41/131 (31.3%) of SLE versus 2/49 (4.1%) of healthy controls were anti-AIM2-positive (p<0.001) (Figure 1A). Interestingly, anti-AIM2 antibodies were strongly associated with both anti-IFI16 and anti-DNA antibodies in the SLE samples measured on the day of visit (Figure 1B and Table 1). We found that anti-AIM2 antibodies were associated with higher measures of SLEDAI (2.29 ± 2.3 versus 1.05 ± 1.61, p=0.0026, Table 1), which was largely driven by the immunology component. Anti-AIM2 antibodies were associated with the presence of disease activity in the skin domain of the SLEDAI index at the date of blood draw: 11/41 (26.8%) of anti-AIM2 positive patients had scores >0 in this domain, compared to 11/90 (12.2%) of anti-AIM2 negative patients (p=0.0463). Anti-AIM2 antibodies were also associated with a small but significant increase (0.63 ± 0.55 versus 0.43 ± 0.51, p = 0.0333) in the SLE Physician Global Disease Activity score, which is based solely on clinical estimation of SLE activity, rather than serologic indices. A multivariable analysis correcting for SLEDAI, anti-dsDNA, and C4 results demonstrated that anti-AIM2 antibodies were significantly associated with anti-IFI16 antibodies with an OR of 3.7 (p=0.007, 95%CI 1.44-9.7). A subset of SLE patients (n=9) demonstrated particularly high levels of anti-AIM2 antibodies with normalized OD > 20 (Figure 1A). These patients had higher SLEDAI values than both lower level anti-AIM2-positive and anti-AIM2-negative patients (Supplemental Table 1). Among all anti-AIM2 positive patients, we found a higher prevalence of positivity for anti–Ro (18/41, 44% vs 19/90, 21%, p=0.0114) and anti–La (10/41, 24% vs 7/90, 8%, p=0.0125) antibodies (Supplemental Table 2).

**Table 1.**
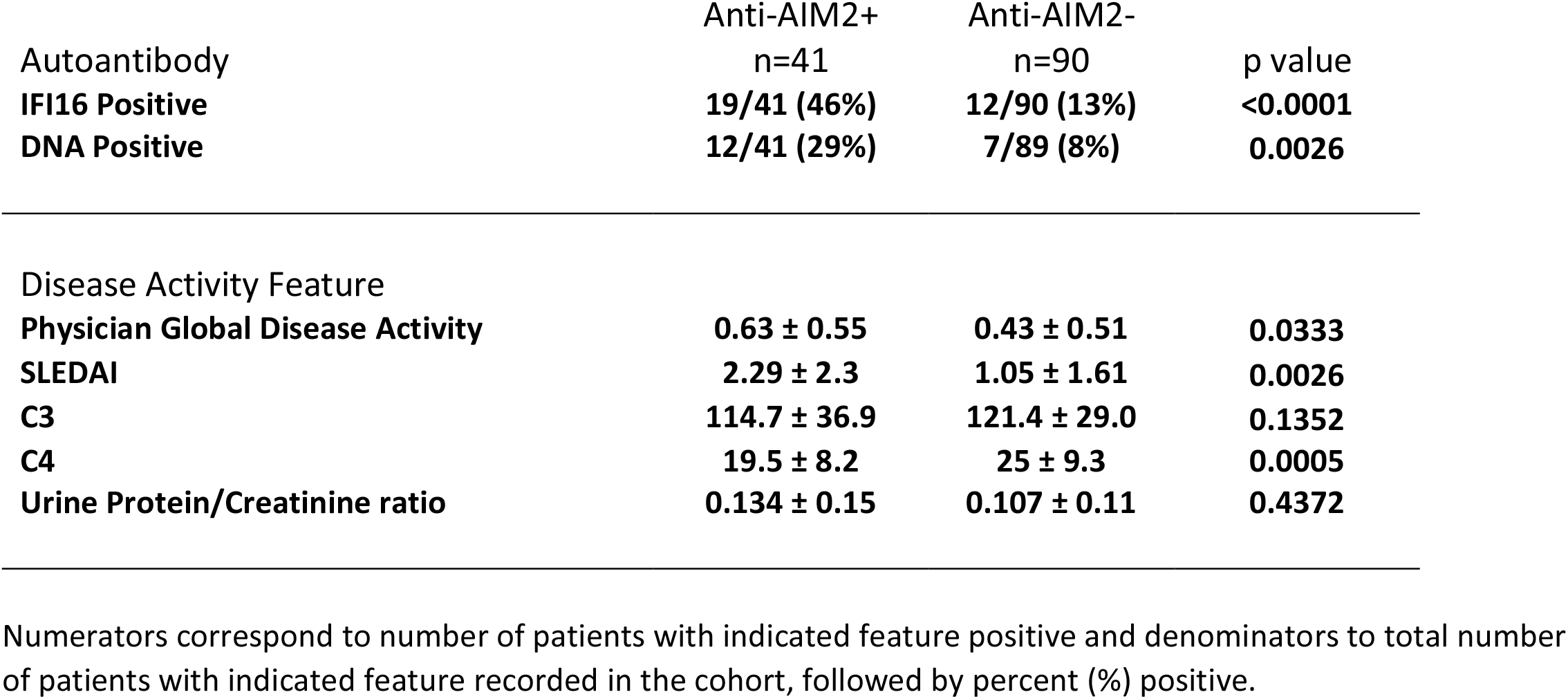
Day of visit phenotypic characteristics of SLE patients related to AIM2 autoantibody status.

| Autoantibody | Anti-AIM2+<br>n=41 | Anti-AIM2-<br>n=90 | p value |
| --- | --- | --- | --- |
| **IFI16 Positive** | **19/41 (46%)** | **12/90 (13%)** | **<0.0001** |
| **DNA Positive** | **12/41 (29%)** | **7/89 (8%)** | **0.0026** |
| Disease Activity Feature |  |  |  |
| **Physician Global Disease Activity** | **0.63 ± 0.55** | **0.43 ± 0.51** | **0.0333** |
| **SLEDAI** | **2.29 ± 2.3** | **1.05 ± 1.61** | **0.0026** |
| **C3** | **114.7 ± 36.9** | **121.4 ± 29.0** | **0.1352** |
| **C4** | **19.5 ± 8.2** | **25 ± 9.3** | **0.0005** |
| **Urine Protein/Creatinine ratio** | **0.134 ± 0.15** | **0.107 ± 0.11** | **0.4372** |

Numerators correspond to number of patients with indicated feature positive and denominators to total number of patients with indicated feature recorded in the cohort, followed by percent (%) positive.

**Figure 1:**
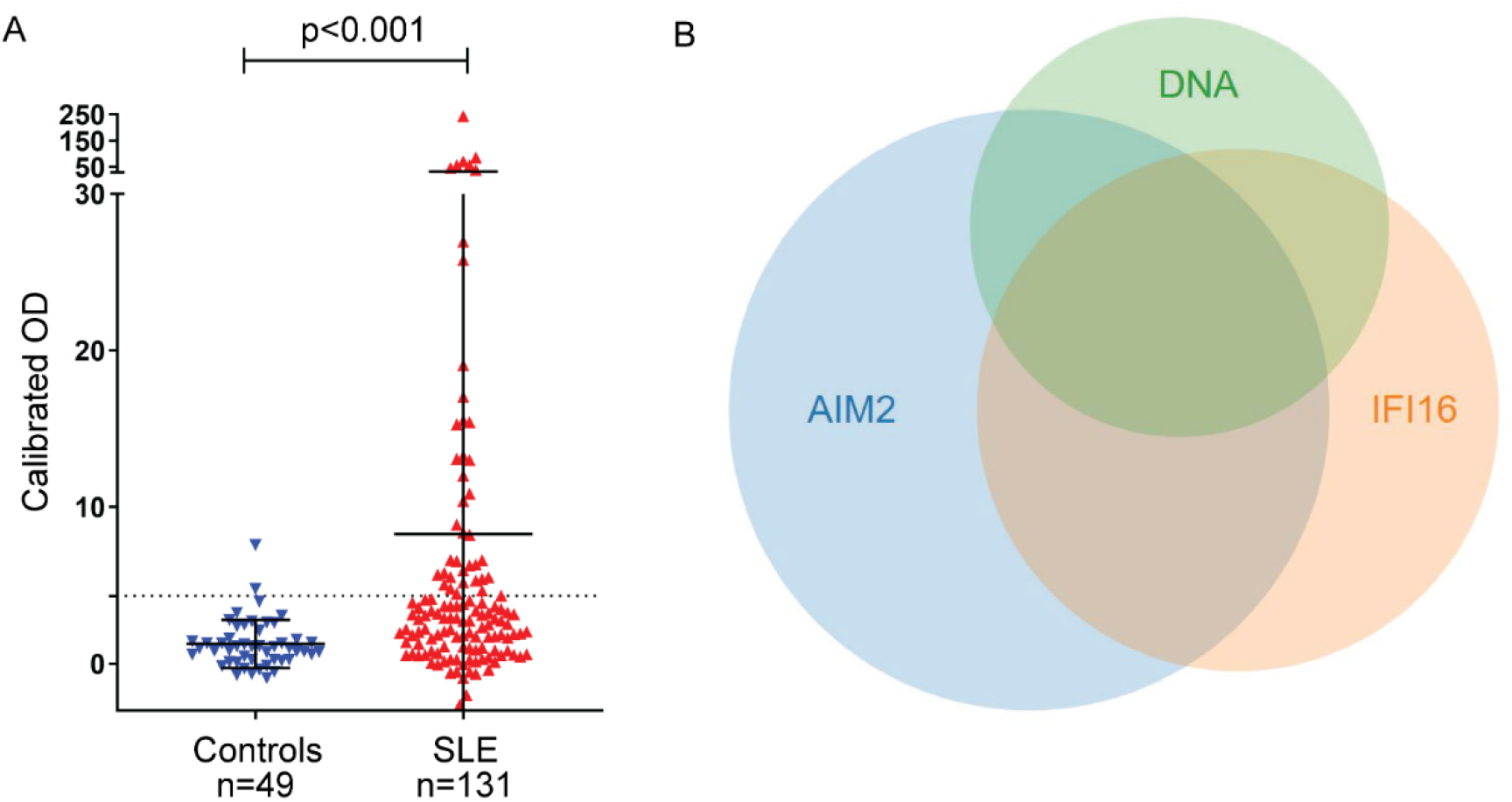
Anti-AIM2 antibodies are associated with anti-IFI16 and anti-DNA antibodies in SLE. AIM2 antibodies were detected using immunoprecipitation of ^35^S-methionine labelled, *in vitro* transcribed and translated protein. Data are presented as OD units calibrated to a known positive reference serum. Dotted line indicates positive threshold value determined as the mean + 2 standard deviations of control serum samples. AIM2 autoantibodies were identified in 2/49 controls and 41/131 SLE patients. Statistical significance was determined using the Mann-Whitney test for nonparametric values (A). Relationship between anti-AIM2, -IFI16 and –DNA antibodies in the SLE cohort (B).

SS shares several phenotypic features with SLE, including the presence of an IFN signature and B cell dysregulation (15), but anti-DNA antibodies are not characteristic of SS. We therefore analyzed SS sera for the presence of anti-AIM2 antibodies, and found 46/133 (34.6%) of SS sera were positive. In contrast to SLE, anti-IFI16 was not enriched in patients with anti-AIM2 antibodies in SS (35% anti-AIM2-positive and anti-IFI16-positive versus 28% anti-AIM2-negative and anti-IFI16-positive in SS, p=0.4324), showing that the association between anti-IFI16 and anti-AIM2 antibodies is specific to SLE, where these immune responses are also associated with anti-dsDNA antibodies.

### AIM2 and IFI16 bind to Neutrophil Extracellular Traps and inhibit their degradation by DNase I

The close relationship between anti-AIM2, anti-IFI16 and anti-dsDNA antibodies in the SLE cohort led us to consider scenarios in which ALR-DNA complexes could be generated and promote the development of autoantibodies against these three antigens. Neutrophil extracellular traps (NETs) have been implicated as important sources of extracellular DNA in SLE, and are linked to the IFN signature as well as autoantibody generation in this disease (16). ALRs are IFN-induced, bind to dsDNA of many origins in a sequence-independent manner, and AIM2 has been identified as a protein constituent of SLE NETs in a proteomics analysis (17). IFI16 is released from epithelial cells undergoing apoptosis (8, 18), and extracellular IFI16 is quantifiable in the sera of SLE patients (11). Considering these observations, we reasoned that when ALRs are generated in the setting of IFN exposure and subsequently released from cells, they might encounter and bind to extracellular NETs, accumulating on this extracellular platform and creating a hub for amplification similar to that observed in the complement and coagulation pathways (19).

To test this hypothesis, we used NETs as a DNA substrate for ALR binding: neutrophils were stimulated to undergo NETosis with PMA, and then incubated with fluorescently labeled IFI16 and AIM2 proteins. We found that both ALRs bind readily to NETs (Figure 2A-B). Co-localization of AIM2 and IFI16 along NET chromatin fibers was visible by confocal microscopy in this analysis, suggesting that both ALRs assemble into filaments on NET DNA.

**Figure 2:**
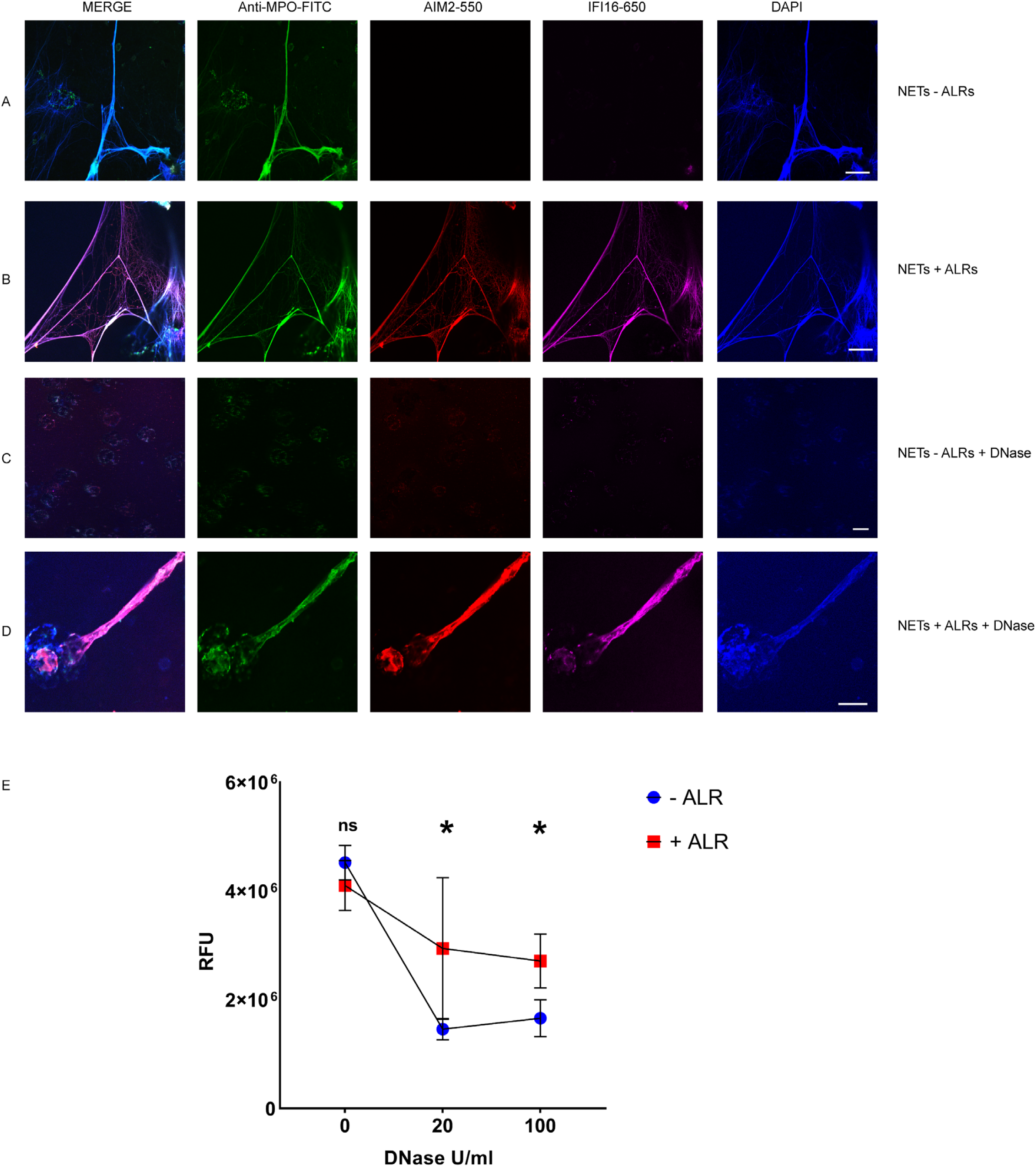
IFI16 and AIM2 bind NETs and prevent NET degradation by DNase I. NETs were induced in neutrophils using PMA 100 nM for 3 hours, then left untreated (A) or incubated with fluorescently labeled AIM2 (pink) and IFI16 (red) at 200 nM at RT for 1 H (B). Following ALR incubation, samples were stained with anti-MPO-FITC antibody (green) and DAPI (blue), then imaged by confocal microscopy. NETs were treated with DNase I at 20 U/mL at RT for 1 hr (C). NETs incubated with ALRs as in (B) were then treated with 20 U/mL DNase I for 1 hr (D). Scale bars = 20 µm. NETs in 96 well plates were incubated with ALRs at 200 nM (or buffer only) for 1 hr at RT, then treated with DNase I at 0, 20, and 100 U/mL for 30 minutes at RT. NETs were then stained with Sytox-Green 5 µM, and samples analyzed by fluorimetry (E). RFU = fluorescence units. Mean and standard deviation of 4 replicate wells are indicated. Mann-Whitney test was used to compare groups. p> 0.05 = not significant (ns). p < 0.05 = significant (*).

IFI16 and AIM2 nucleoprotein filaments are highly stable and persist even after the dsDNA template has been degraded by nucleases (8, 20). We therefore hypothesized that ALR-bound NETs might resist nuclease exposure, potentially enhancing antigenicity. We used DNase I to explore this question, as DNase I is the nuclease responsible for effective clearance of NETs (21), and DNase I deficiency has been associated with SLE in both human subjects and animal models (22). After exposure to 20 U/ml DNase I for 1 hour at RT, both myeloperoxidase (MPO) and DNA signals were completely degraded, leaving no observable fluorescence in any channel (Figure 2C). When NETs were first incubated with ALRs, however, we observed incomplete ALR-NET degradation by DNase I – in some areas, IFI16 and AIM2 remained present and co-localized with MPO (Figure 2D). In addition, there was observable DNA remaining in these areas of persistent ALR structures, implying that the ALRs had partially shielded NET DNA from degradation. This finding suggested that both the protein and DNA components of the ALR-NET structure are resistant to DNase-mediated clearance. To better quantify this, we employed a plate-based Sytox Green assay to measure the dsDNA content of NETs following exposure to DNase I (Figure 2E). This assay confirmed that ALR-bound NETs are resistant to DNase I, leaving more DNA present following nuclease treatment (Figure 2E). Together these experiments demonstrate that ALRs bind to NETs, generating a protein-DNA structure with enhanced resistance to DNase-mediated clearance.

### IFI16-NETs are present in lupus nephritis

Prior studies have presented evidence of in vivo NET formation within the renal tissues of SLE patients, supporting the notion that dysregulated neutrophil function contributes to immune pathology in this disease (23). We therefore sought to determine whether ALR-NET interactions could be identified among NETs present in lupus nephritis biopsies. Considering that patients with diffuse proliferative lupus nephritis are known to harbor netting neutrophils in renal tissue (23), we identified patients with diffuse proliferative lupus nephritis, then selected 5 samples whose biopsies demonstrated neutrophilic infiltrates or karyorrhectic debris (Supplemental Table 3). We found that AIM2 was highly expressed in MPO-positive infiltrating cells (Figure 3A), while IFI16 was expressed more broadly throughout renal cell types (Figure 3B). We detected NETs containing both AIM2 and IFI16 in glomerular and interstitial infiltrates (Figure 3C and D). High magnification, z-stack imaging (Supplemental Figure 1) confirmed that these structures represented extracellular DNA that co-stained for MPO and AIM2 or IFI16, consistent with ALR-bound NETs, rather than adjacent or overlapping cell nuclei. In summary, our immunostaining experiments provide evidence that AIM2 and IFI16 bind NETs in the setting of diffuse proliferative lupus nephritis, establishing AIM2 and IFI16 as NET-bound SLE autoantigens.

**Figure 3:**
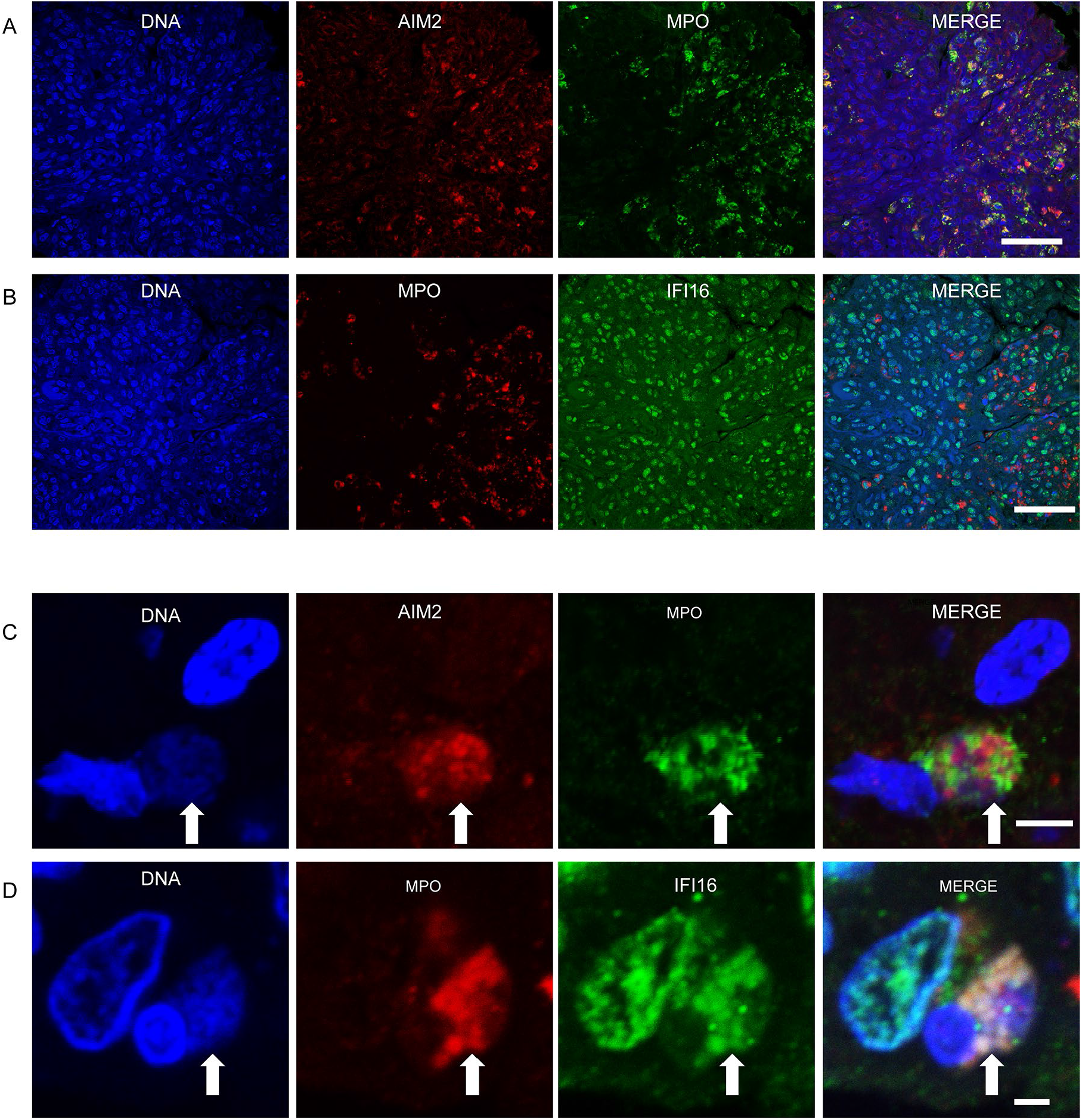
IFI16 and AIM2 bind NETs in diffuse proliferative lupus nephritis. Representative images of ALR expression and ALR-NETs identified in patients with class IV lupus nephritis. AIM2 (A) expression was largely detected in MPO expressing neutrophils, while IFI16 (B) was more broadly distributed. NETs (arrows) demonstrating co-localizing staining for DNA, MPO, and AIM2 (C) or IFI16 (D) visualized by confocal microscopy. Scale bars: 50 µm (A, B) 5 µm (C), 2 µm (D).

## Discussion

SLE features autoantibodies that bind nucleic acids and nucleic acid-binding proteins, and extracellular nucleic acids contribute to SLE pathogenesis (24). Here, we identify the dsDNA sensor AIM2 as a novel autoantigen in SLE, and demonstrate that anti-AIM2 antibodies are associated with SLE disease activity markers. Furthermore, we find that NETs provide a scaffold for ALR oligomerization, which in turn confers resistance to nuclease degradation.

NETosis is a process whereby dsDNA is expelled into the extracellular space at sites of tissue damage, and is of mechanistic relevance in SLE (16). The NET dsDNA scaffold is a structure on which a variety of molecules can interact, and is a source of antigenic proteins in SLE and other inflammatory diseases (12, 25). We found that both IFI16 and AIM2 readily assemble into filaments along the length of NET dsDNA. Unexpectedly, we found this ALR-NET structure resists DNase-mediated degradation. NETs promote IFN signaling at sites of their generation when engulfed by immune cells (26, 27), and may have additional disease-amplifying functions (28). By prolonging the stability of interferogenic NETs, extracellular ALRs may enhance IFN signaling at sites of neutrophil activation, which could be further amplified by IFN-induced expression of the ALRs themselves.

Impairment of NET removal has been specifically linked to the presence of lupus nephritis (21) - a manifestation of SLE with significant associated morbidity (29). Neutrophilic infiltration of the kidney is a feature of more severe forms of glomerulonephritis, and NET formation in this organ may contribute to renal damage through the propagation of IFN signaling, immune cell activation and thrombosis (28). Confocal microscopy has been utilized to demonstrate the presence of NET structures in renal lesions of patients with SLE (23, 30) and also ANCA associated vasculitis (31), supporting the notion that NETs play a pathogenic role in the immune dysregulation and tissue damage that occur in glomerulonephritis.

Here, we demonstrate for the first time that the DNA sensors AIM2 and IFI16 bind to NETs in vivo, through imaging studies of proliferative lupus nephritis specimens. Our data include z-stack images at high magnification, clearly demonstrating the presence of extracellular DNA-MPO-ALR complexes in this site. This finding supports previous data (11, 32, 33) suggesting that the ALRs may have important functions not just intracellularly, but also in the extracellular environment. The large chromatin fibers generated through NETosis represent sizeable dsDNA templates upon which IFI16 and AIM2 monomers oligomerize in the extracellular space, and are expected to result in durable immunostimulatory structures at sites of IFN-induced protein expression. ALR-bound NETs therefore may promote not only local immune activation, but the targeting of ALRs (and DNA) by antibodies in SLE.

Our data indicate that AIM2 is targeted not only in SLE but also in SS – a condition in which NETosis has not been linked to disease pathology. In contrast to the relationship seen in SLE, we found no association between AIM2 and IFI16 antibodies in SS, and anti-dsDNA antibodies are absent in SS. This difference highlights the important role of disease-specific tissue processes in the development of unique autoantibody profiles against shared antigens. In the setting of lupus nephritis, we observed ALR expression by both neutrophils and resident renal cells, and suspect that these antigens may be released by a variety of cell types in the kidney, leading to the observed extracellular interaction with NET DNA. Contrastingly, neutrophil infiltration in target salivary tissues is not a common feature of SS, and the absence of the NET DNA scaffold may explain the differing autoantibody profile observed in that condition.

In summary, we have identified the ALRs AIM2 and IFI16 as NET-binding autoantigens in SLE. The ALR-NET interaction may increase NET longevity and perpetuate NET-mediated inflammatory signaling in lupus nephritis and other sites of NET generation and IFN expression. This work supports a role for the ALRs in extracellular immune processes as NET-binding antigens in SLE.

---

**Supplemental Figure 1:**
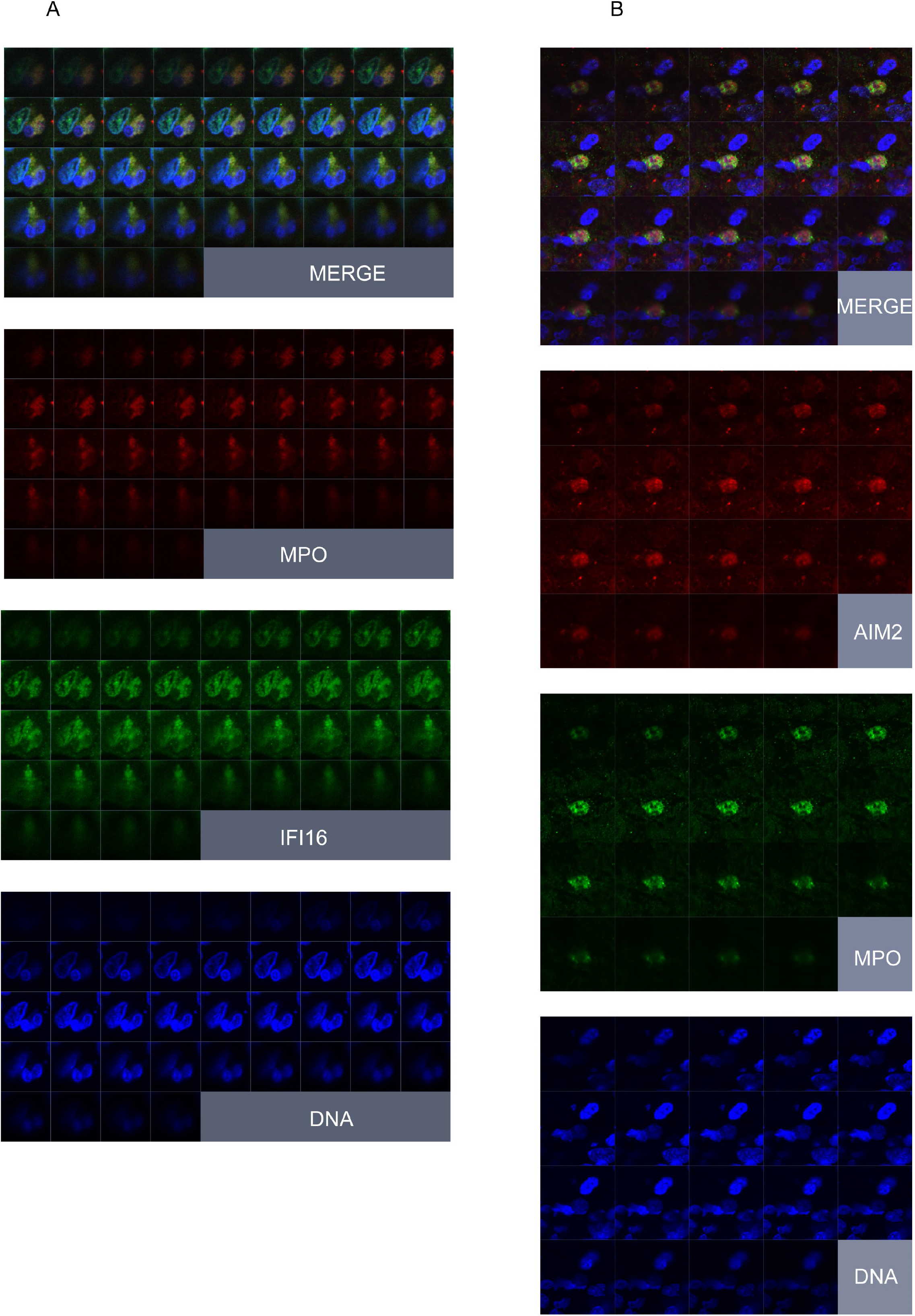
Z-stack imaging of AIM2/IFI16-NETs in lupus nephritis. Renal biopsy paraffin section stained for DNA, MPO and IFI16/AIM2 and imaged using z-stacking to identify extracellular DNA-containing structures containing MPO and IFI16 or AIM2. Individual squares within each panel represent adjacent focal planes, proceeding sequentially from top left to bottom right in each area imaged.

**Supplemental Table 1.** Phenotypic Characteristics of SLE Patients Related to AIM2 Autoantibody Level.

| Feature | (1) Anti-AIM2+<br>**OD> 20**<br>n=9 | (2) Anti-AIM2+<br>**4.3<OD<20**<br>n=32 | (3) Anti-AIM2-<br>**OD<4.3**<br>n=90 | p value<br>1 vs 2 | p value<br>1 vs 3 | p value 2<br>vs 3 |
| --- | --- | --- | --- | --- | --- | --- |
| Age (years) at blood draw, mean ± SD | 46.7 ± 13.2 | 52.8 ± 11.4 | 52.1 ± 13.8 | 0.1431 | 0.2338 | 0.7120 |
| Physician Global Disease Activity | 0.73 ± 0.28 | 0.61 ± 0.60 | 0.44 ± 0.51 | 0.2735 | **0.0156** | 0.2014 |
| SLEDAI | 3.77 ± 2.11 | 1.88 ± 2.21 | 1.05 ± 1.61 | **0.0249** | **0.0001** | 0.0742 |
| IFI16 Positive | 7/9 (78%) | 12/32 (38%) | 12/90 (13%) | 0.0570 | **<0.0001** | **0.0080** |
| DNA positive | 6/9 (67%) | 6/32 (19%) | 8/90 (9%) | **0.0106** | **0.0002** | 0.1929 |
| C3 (mg/dL) | 104.1 ± 29.0 | 117.7 ± 38.7 | 121.4 ± 29.0 | 0.2204 | 0.1069 | 0.3291 |
| C4 (mg/dL) | 14.9 ± 7.7 | 20.8 ± 8.0 | 25.0 ± 9.3 | **0.0378** | **0.0019** | **0.0084** |

Numerators correspond to number of patients with indicated feature positive and denominators to total number of patients with indicated feature recorded in the cohort, followed by percent (%) positive.

**Supplemental Table 2.**
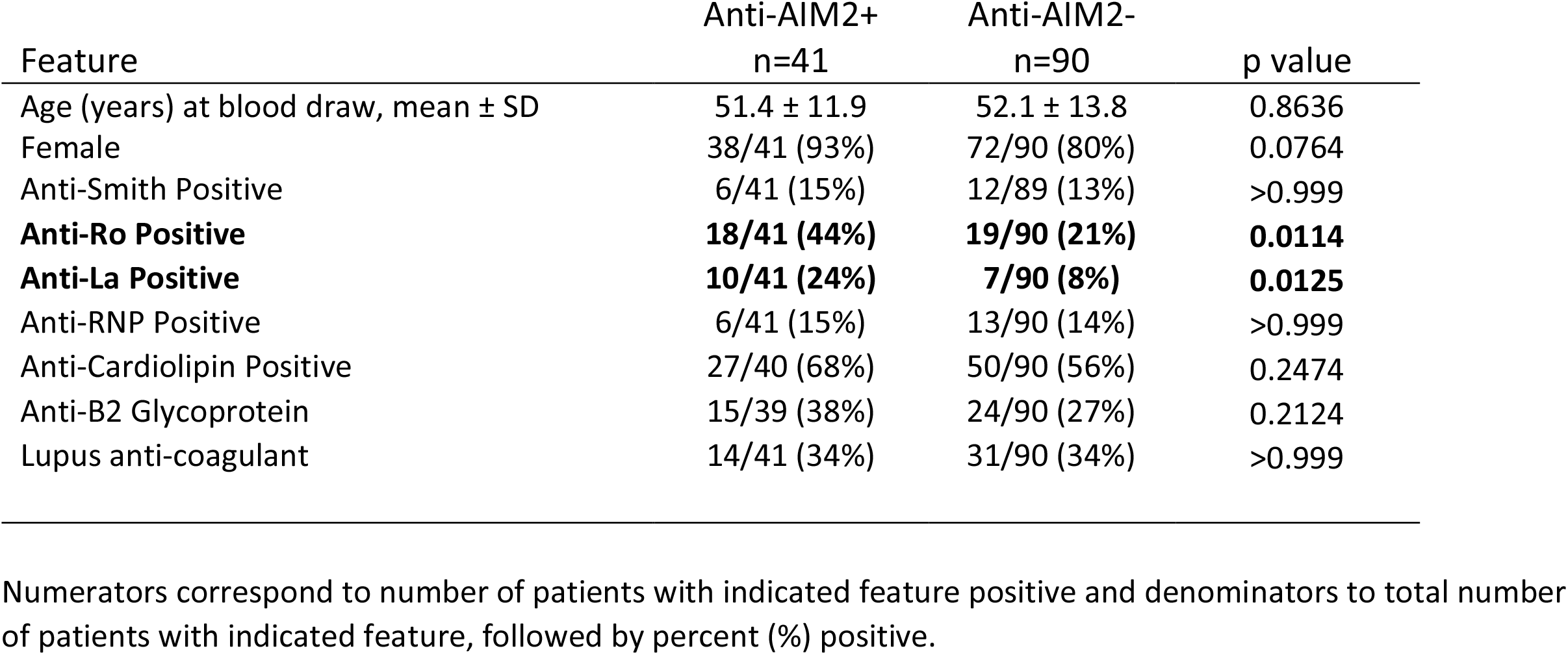
Immunologic phenotype of SLE patients related to AIM2 autoantibody status.

**Supplemental Table 3:** Lupus nephritis renal biopsies used in confocal imaging.

| <u>Age</u> | <u>Sex</u> | <u>Diagnosis</u> |
| --- | --- | --- |
| 36.9 | F | Diffuse proliferative lupus nephritis, ISN/RPA Class IV-A/C (S) with early segmental consolidation. |
| 36.7 | F | Diffuse proliferative and membranous lupus nephritis with focal segmental glomerulosclerosis (ISN/RPS CLASS IV A/C-G + V) |
| 19.9 | F | Diffuse proliferative and focally necrotizing lupus glomerulonephritis, ISN/RPS class IV-G (A) and tubulointerstitial inflammation with scattered interstitial immune complex deposits. |
| 12.9 | F | Diffuse proliferative lupus nephritis, ISN/RPS class IV-G (A/C) |
| 28.6 | F | Diffuse proliferative lupus nephritis; ISN/RPS class IV-G (A) |

